# Metatranscriptome of infected root canals in teeth with apical periodontitis

**DOI:** 10.1101/2022.12.08.519614

**Authors:** Ericka T. Pinheiro, Giancarlo Russo, George T. M. Candeiro, Bruna G. Vilela, Brenda P. F. A. Gomes, Thomas Attin, Lamprini Karygianni, Thomas Thurnheer

## Abstract

Metatranscriptomics has not yet been applied to the study of the endodontic microbiome. Therefore, this study investigated the microbial composition and patterns of gene expression in teeth with asymptomatic apical periodontitis. We performed next-generation, microbiome-wide RNA-sequencing in eight root canal samples, including five primary and three secondary endodontic infections. Proteobacteria, Bacteroidetes, Firmicutes and Actinobacteria represented the dominant phyla, whereas Fusobacteria, Spirochaetes and Synergistetes were among the non-dominant phyla. At species level, the microbiome was dominated by Gram-negative and Gram-positive anaerobes. In particular, *Prevotella* spp. and *Olsenella uli* were more abundant in primary than in secondary infections. Transcripts encoding moonlighting proteins were abundant, including glycolytic proteins, translational elongation factors, chaperonin, and heat shock proteins. *Fusobacterium* was highly active in primary infections, showing upregulation of genes involved in protein synthesis, responses to oxidative stress and resistance to antibiotics, especially tetracycline. In addition, other members of the community expressed high numbers of antimicrobial resistance genes. Overall, metatranscriptomics revealed the expression of putative virulence factors and the activity of potential endodontic pathogens, represented mainly by obligate anaerobes. In particular, the functional role of *Fusobacterium* in primary infections was highlighted. Furthermore, antimicrobial resistance, especially to tetracycline, was a common feature in endodontic samples.

## 1. Introduction

Apical periodontitis is an inflammatory disease of the apical periodontium caused primarily by an infection in the root canal of a tooth, typically associated with pulp necrosis (primary endodontic infections) [1,2]. Failure of endodontic therapy to definitively clear the infection or prevent reinfection can lead to persistent/secondary infections, characterized by the persistence of apical periodontitis [1,2]. Over the last decades, high-throughput sequencing has enabled the screening of the 16S ribosomal RNA (rRNA) region in hundreds of thousands of microbial ecosystems, including the healthy and diseased oral microbiome [3], therefore also expanding the knowledge of the bacterial communities in infected root canals [4-12]. In particular, a recent study based on the combined analysis of 16S rRNA and cDNA of infected root canal microbiomes showed that only part of community is transcriptionally active [12]. Moreover, the aforementioned study showed that uncultivated/ difficult-to-culture bacteria were active members in the endodontic microbial community. However, as this study was restricted to the analysis of 16S rRNA, no additional information about microbial interactions or virulence was revealed.

In recent years, metatranscriptomics (*i*.*e*., the analysis of collective gene expression) has provided insights into how microbial communities behave in oral health and disease. Thus far, studies on the metatranscriptome of the oral microbiome have been restricted to 2 diseases: caries and periodontal disease [13-21]. Indeed, the knowledge on the sugar metabolism in caries was expanded by metatranscriptomics [18]. In the case of periodontal disease, this approach revealed that microbial communities displayed conserved metabolic gene expression profiles despite their diverse composition [16,20]. Moreover, metatranscriptomics revealed “keystone” pathogens in periodontal disease [21].

To the best of our knowledge, metatranscriptomics has not yet been applied to the functional analysis of the endodontic microbiome. Therefore, the aim of this study was to investigate the gene expression of active players in the endodontic microbiome of teeth with asymptomatic apical periodontitis apical periodontitis, including either primary or persistent/secondary endodontic infections. The metatranscriptomic analysis may cast some new light on the nature of endodontic pathogens and on the core activities associated with apical periodontitis.

## 2. Results

### 2.1. Overview of the microbial community

First, we analyzed the composition of the microbial community in the dataset. The overall variability was rather high across samples, with a total of 166 species identified in eight samples, belonging to ten different phyla [Supp. Table 1]. The average number of identified species across the samples was 121, with EP08 being the least populated sample (95 species) while EP10 harbored 141 different species [Supp. Table 2]. None of the species identified were present exclusively in one sample, while 49% (78) were present in all samples [Figure 1a]. In particular, 103 species were common to all samples with primary infections, and 79 to all samples with secondary infections [Figure 1b,1c].

**Figure 1.**
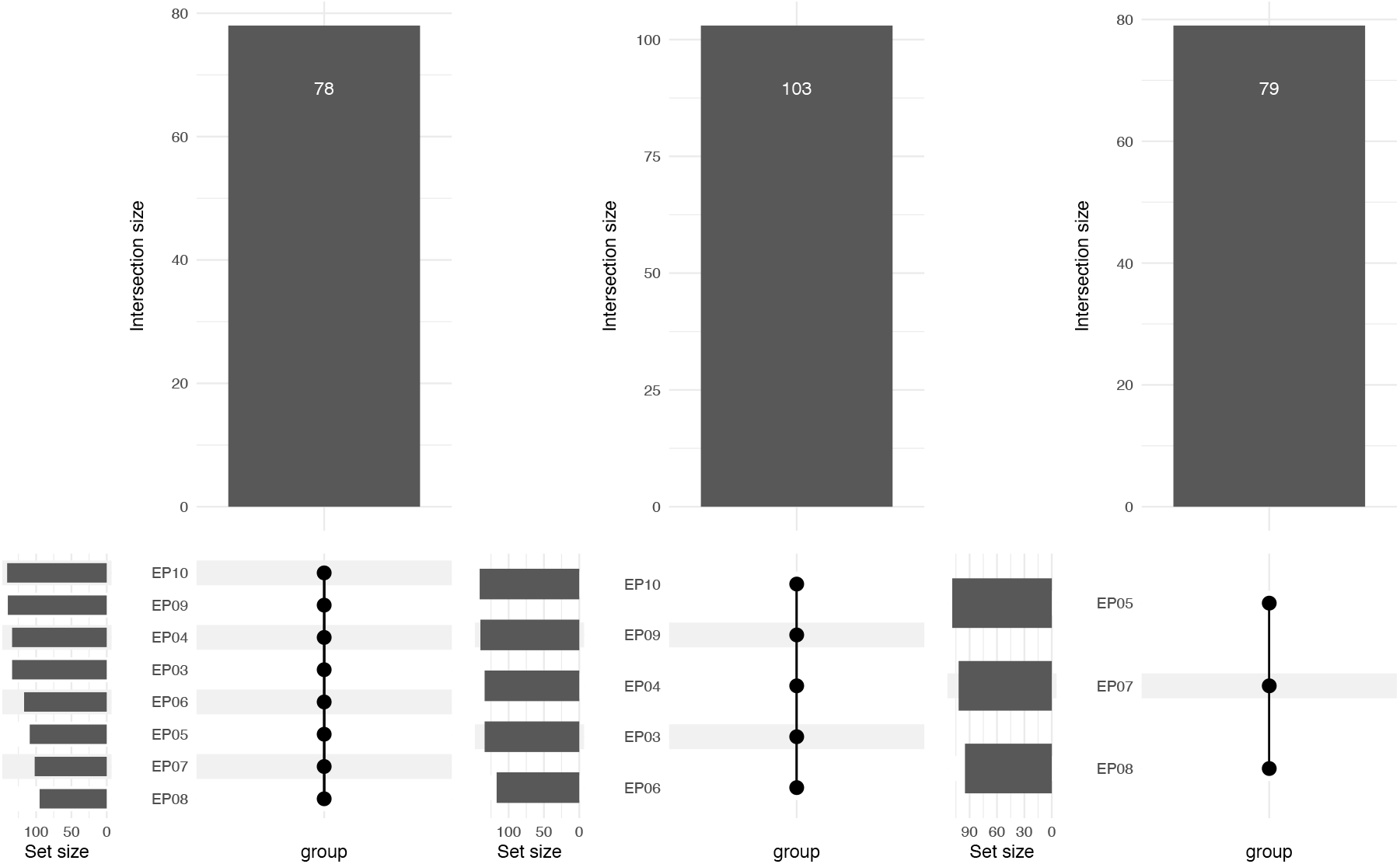
Number of species present in (a) all root canal samples, (b) in samples from primary endodontic infections (EP03, EP04, EP06, EP09, EP10) and (c) in samples from secondary endodontic infections (EP05, EP07, EP08).

Of the 166 total species, 53, belonging to nine different phyla, represented the most abundant taxonomic groups (Table 1). Altogether, Proteobacteria (28.06%), Bacteroidetes (26.70%), Firmicutes (19.68%) and Actinobacteria (19.58%) constituted the core microbiome of the most abundant phyla by about an order of magnitude. Fusobacteria (2.68%), Synergistetes (1.88%) and Spirochaetes (0.67%) were identified in at least five samples and represented more than 0.5% of the community, suggesting the presence of a secondary core group of phyla. Despite the presence of core organisms, the sample-wide variance in their abundance was high [Figure 2a]. In some cases, the most prevalent phyla shared comparable levels of relative abundance (e.g., EP07, EP04), while in other cases one phylum was clearly dominating (e.g., EP03, EP08, EP09) [Figure 2a]. The ten most abundant genera and species are represented in Figure 2, panels b and c, respectively.

**Table 1.** The most abundant phyla and genera in root canal samples.

| Actinobacteria | Bacteroidetes | Elusimicrobia | Firmicutes | Fusobacteria | Gemmatimonadetes | Proteobacteria | Spirochaetes | Tenericutes |
| --- | --- | --- | --- | --- | --- | --- | --- | --- |
| *Actinomyces* | *Bacteroides* | *Eubacterium* | *Bacillus* | *Fusobacterium* | *Gemmatirosa* | *Desulfovibrio* | *Treponema* | *Mycoplasma* |
| *Atopobium* | *Capnocytophaga* |  | *Clostridium* |  |  | *Enterobacter* |  |  |
| *Cutibacterium* |  |  | *Filifactor* |  |  | *Haemophilus* |  |  |
| *Olsenella* | *Prevotella* |  | *Mogibacterium* |  |  | *Klebsiella* |  |  |
| *Propionibacterium* | *Tannerella* |  | *Parvimonas* |  |  | *Neisseria* |  |  |
| *Rothia* |  |  | *Selenomonas* |  |  | *Pasteurella* |  |  |
|  |  |  | *Staphylococcus* |  |  | *Pseudomonas* |  |  |
|  |  |  | *Streptococcus* |  |  | *Ralstonia* |  |  |

**Table 2.** The 10 species with at least 10 AMR homologues. They are sorted by the ratio between the number of AMR genes and total genes.

| AMR | Total Genes | Ratio | Species |
| --- | --- | --- | --- |
| 21 | 320 | 0.06 | Tannerella forsythia |
| 17 | 265 | 0.06 | Capnocytophaga sp. oral taxon 323 |
| 23 | 625 | 0.04 | Olsenella uli |
| 71 | 2096 | 0.03 | Cutibacterium acnes |
| 29 | 1041 | 0.03 | Staphylococcus epidermidis |
| 23 | 796 | 0.03 | Parvimonas micra |
| 20 | 562 | 0.03 | Prevotella intermedia |
| 10 | 361 | 0.03 | Fusobacterium nucleatum |

**Figure 2.**
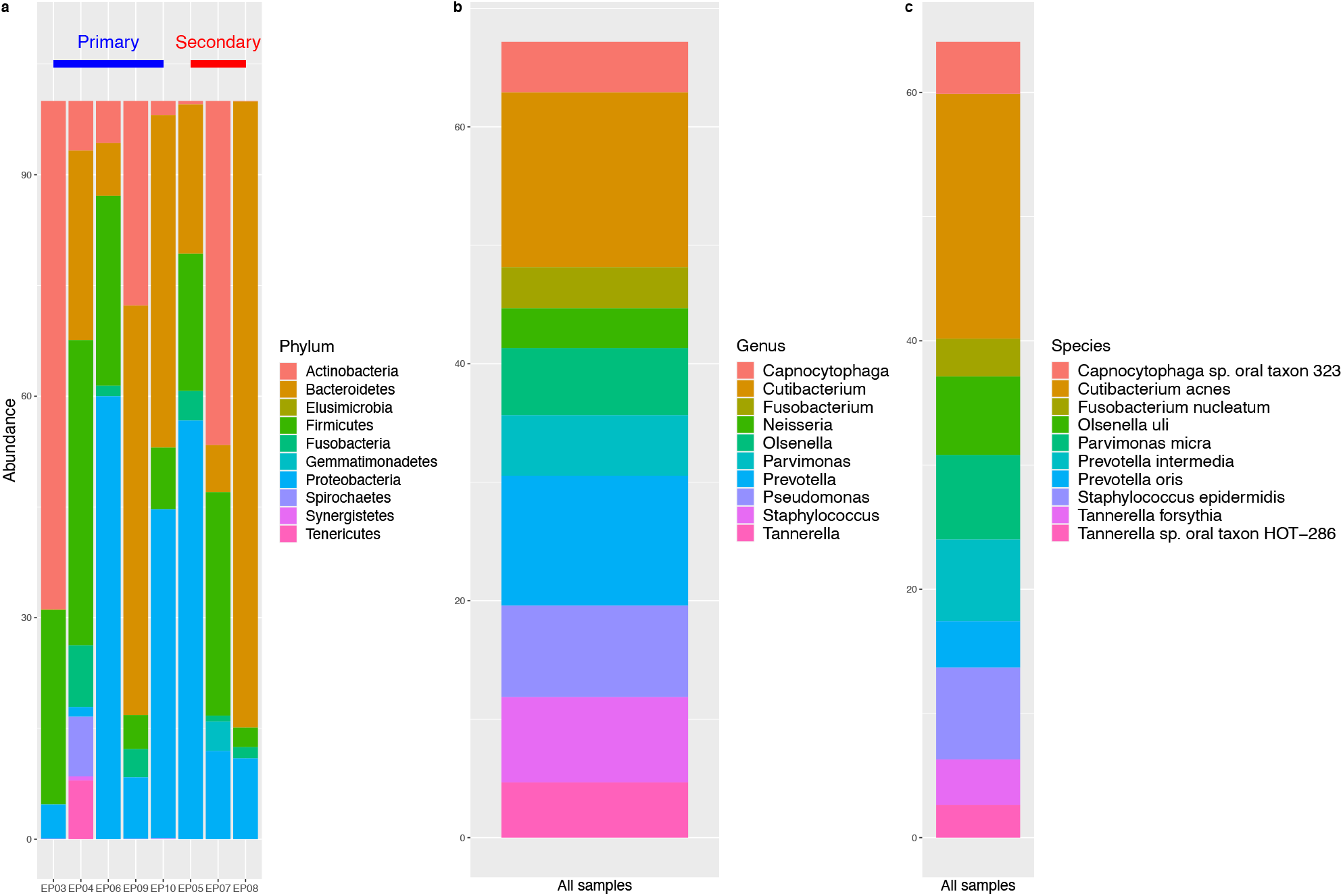
Relative abundance (%) of bacterial phyla in root canal samples of 5 teeth with primary endodontic infections and 3 with secondary endodontic infections (a) and the ten most abundant genera (b) and species (c).

We then statistically tested the difference in microorganism’s abundance between primary and secondary infections. Using a conventional threshold of 0.05 for the p-value adjusted for multiple testing, seven species resulted significantly differentially abundant [Figure 3a]. *Olsenella uli* (p = 2.0e-04) showed the highest discrepancy, being among the top ten most abundant species in primary infections and completely depleted in secondary infections. An additional five species, which included four *Prevotella* spp. (*P. denticola, P. dentalis, P. enoeca*, and *P. intermedia*) were also more abundant in primary infections, and depleted in secondary infections. The only species significantly more abundant in secondary infections was *Bacillus cereus* (p = 7.3e-02).

**Figure 3.**
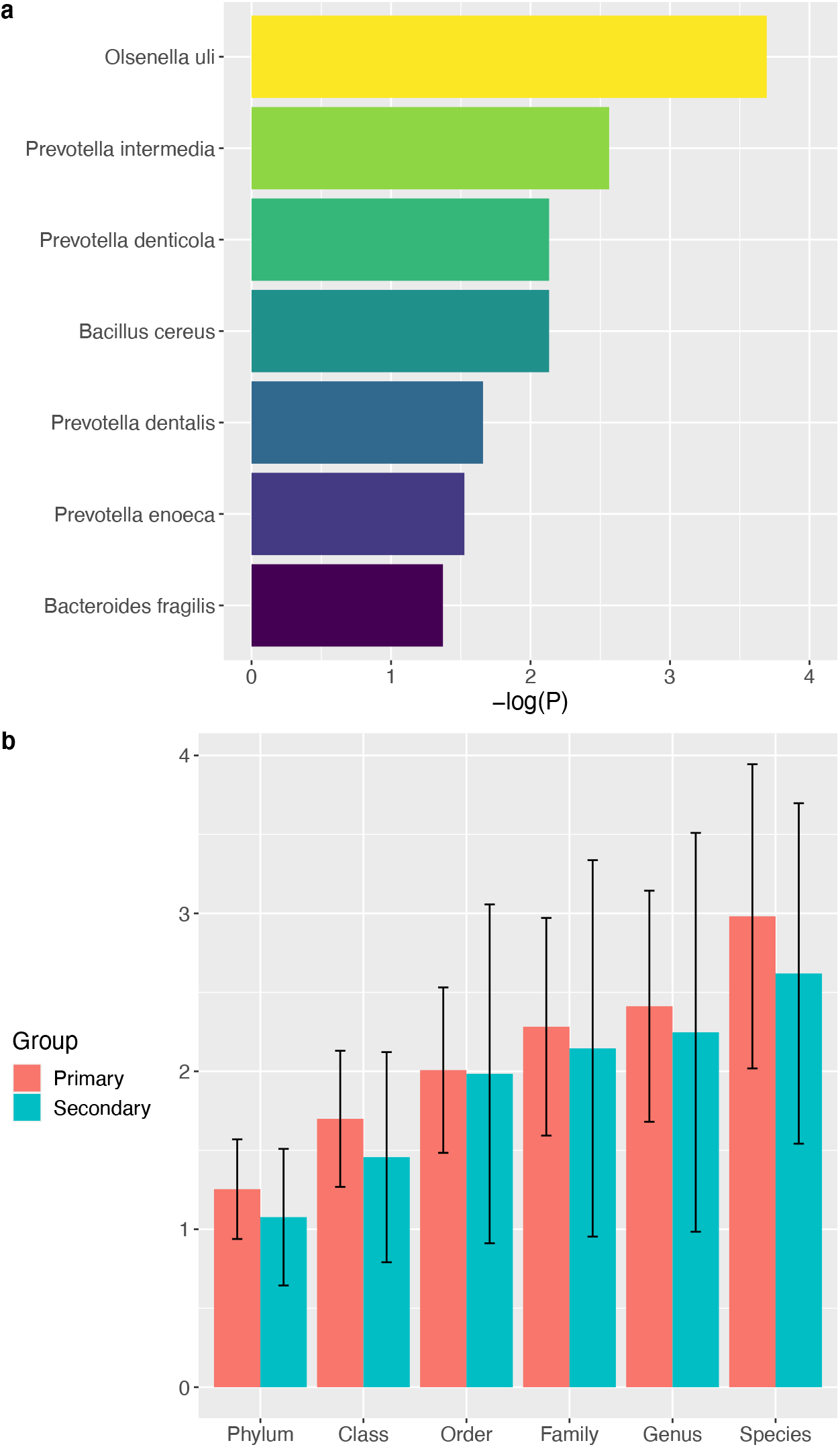
(a) Species showing a significant differential abundance between the primary and the secondary groups (adjusted p-threshold = 0.05) (b) Bacterial diversity, as measured by the Shannon index, in primary and secondary endodontic infections.

Finally, in each sample the alpha diversity was estimated using the Shannon index. The microbiomes from secondary infections presented slightly lower diversity at all taxonomic ranks [Figure 3b]. However, none of these differences was statistically significant.

### 2.2. Summary of the metatranscriptomes

The assembly of the metatranscriptomes revealed 152 species transcriptionally active in at least 2 samples, fostering a number of expressed putative transcripts between 3 (*Arthrobacter* sp. DCT-5) and 4061 (*Cutibacterium acnes*), with an average of 148. Out of those 78 species present in all samples, 71 (91 %) have at least one gene shared by all samples.

The average absolute number of putative transcripts which were annotated through matching a Uniprot ID was 72, ranging from 0 (five species) to 2096 (*Cutibacterium acnes*), and the percentages of annotated genes for each species, relative to the total number of transcripts in the same species, varied from 0% to 100% (five species) [Figure 4a]. In about half of the species with at least one Uniprot annotation (72/147), different putative transcripts were annotated to the same Uniprot ID. However, such one-to-many transcript-Uniprot ID relationships represent the exception: on average across the species with at least one case of identical annotation, 87.56% of the transcripts were still annotated to only one Uniprot ID [Figure 4a]. In total, across the metatranscriptomes of the eight samples analyzed, we identified 10,928 annotated transcripts, corresponding to 6,514 unique Uniprot IDs, which in turn mapped to 3,084 unique gene IDs.

**Figure 4.**
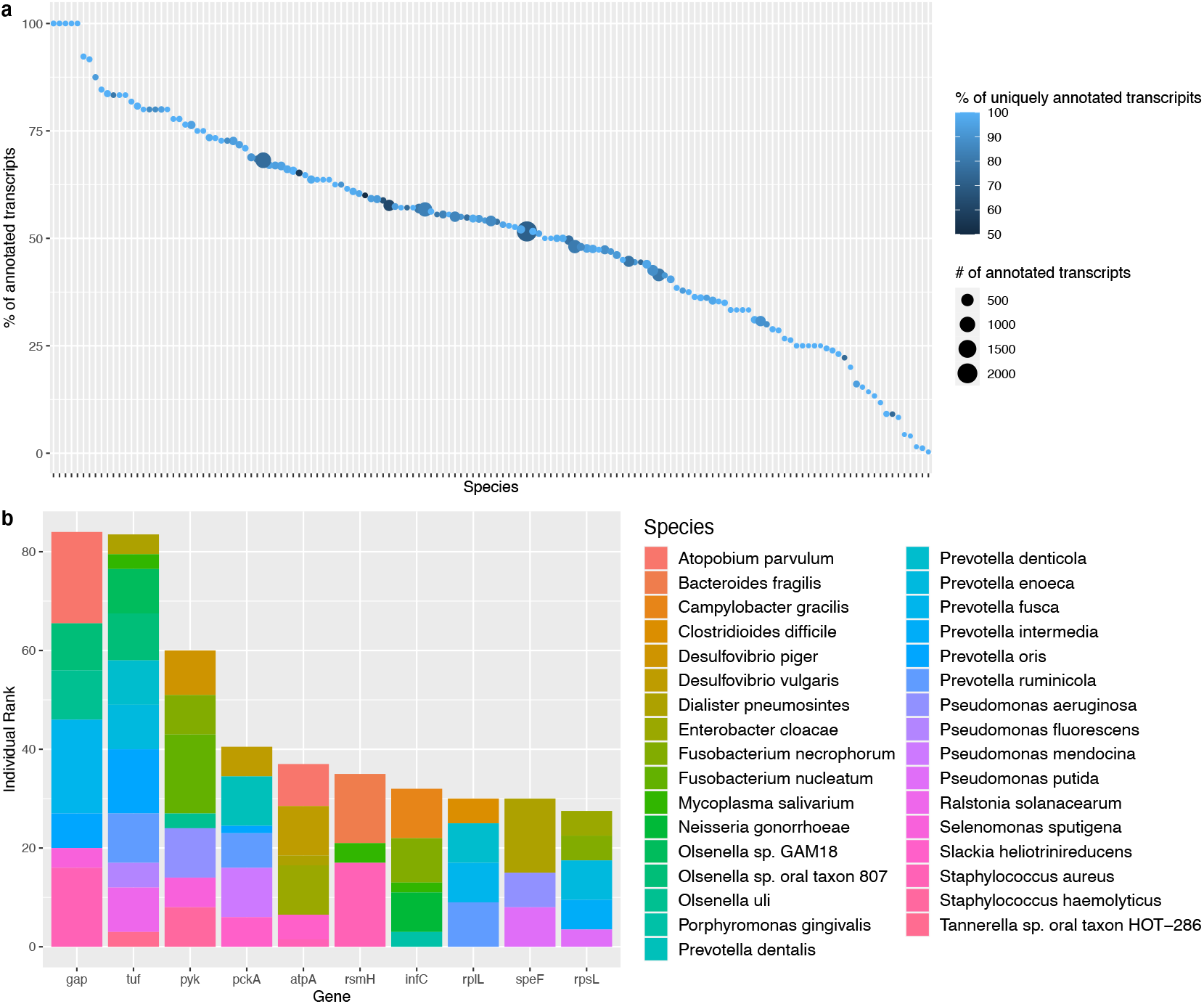
(a) Percentages of annotated genes for each species (relative to the total number of transcripts in the same species); (b) the top 10 most abundant annotated transcripts in each species.

To gauge which species showed the highest degree of transcriptional activity across the whole transcriptome, we calculated, for the species present in all individuals, a normalized coefficient of variation (NCV) of expression levels of the transcripts averaged across the samples. Briefly, in each species, we calculated the coefficient of variation (mean/sd) of each transcripts across the samples. We then scaled those by the fraction of samples with non-zero counts and then took the averages across the transcripts. We restricted the calculations to the annotated products and detected that *Streptococcus salivarius* yields the highest extent of transcription [Suppl. Table 3].

Among the core species, only in about 13% of the cases (10/78) the transcript with the absolute overall highest expression across the eight samples could be annotated [Suppl. Table 4]. In particular, *pck*A (Phosphoenolpyruvate carboxykinase) was the only gene with the highest expression in more than one species (*Prevotella dentalis* and *Pseudomonas mendocina*). In order to test whether a gene has the highest level of expression in more than one species, we had to restrict to the data from the annotated genes, six of which were associated with the highest level of expression in more than one species.

In particular, *gap* (Glyceraldehyde-3-phosphate dehydrogenase) and *grd*E (glycine reductase) were the most ubiquitous high-abundance gene, covering four species [Suppl. Table 5]. Finally, we wanted to extrapolate the genes with the overall most prominent presence across the species. Therefore, we ranked the top 10 most abundant (on average) annotated transcripts in each species and summed the rank across the species [Figure 4b]. Among the overall most expressed genes, the top 3 were *gap* (Glyceraldehyde-3-phosphate dehydrogenase), *tuf* (elongation factor) and *pyk* (pyruvate kinase), with an overall expression rank 2 to 3 times (*gap* and *tuf*) and 1.5 to 2 times (*pyk*) larger than the average of the genes ranked (3-10).

In recapitulating those numbers, we excluded those genes with an eukaryotic annotation, since these are most likely the result of computational artefacts due to sufficient sequence homology. It is important to remark in this context that it is common that the annotated protein is assigned in Uniprot to a different bacterium from the one in which the gene sequence has been assembled. This is in part due to extremely high level of homology across bacterial species, and in part because in the Uniprot database any given protein is associated to only one organism.

### 2.3. Functional annotation

In order to perform functional annotation, we removed transcripts without a corresponding Uniprot ID, which resulted in the exclusion of the 5 species with no Uniprot annotated transcript (*Arthrobacter* sp.DCT-5, *Comamonas kerstersii, Gemmatirosa kalamazoonesis, Paracoccus mutanolyticus, Spirosoma pollinicola*). For the remaining 147 species, we retained, in those transcripts with a multiple Uniprot annotation, the hit with the lowest blastx p-value. In 41.5% of the species (61/147), each protein ID was associated with a different protein family, while in the remaining 58.5% several proteins belonged to the same family. In total, 1,612 protein families were associated with the annotated Uniprot IDs. The overall predominant protein family was the ATP-binding cassette (ABC) transporter superfamily, to which 163 Uniprot ID (1.94%) are assigned. The 20 most abundant protein families are listed in Figure 5a. They mostly include GTPase-, RNA polymerase- and ATP/pyruvate families, which are typically involved in the basic molecular mechanisms. Additionally, transcripts encoding stress response proteins were abundant in root canal samples, including pyruvate:ferredoxin/flavodoxin oxidoreductase-, chaperonin (HSP60)- and heat shock protein 70 families.

**Figure 5.**
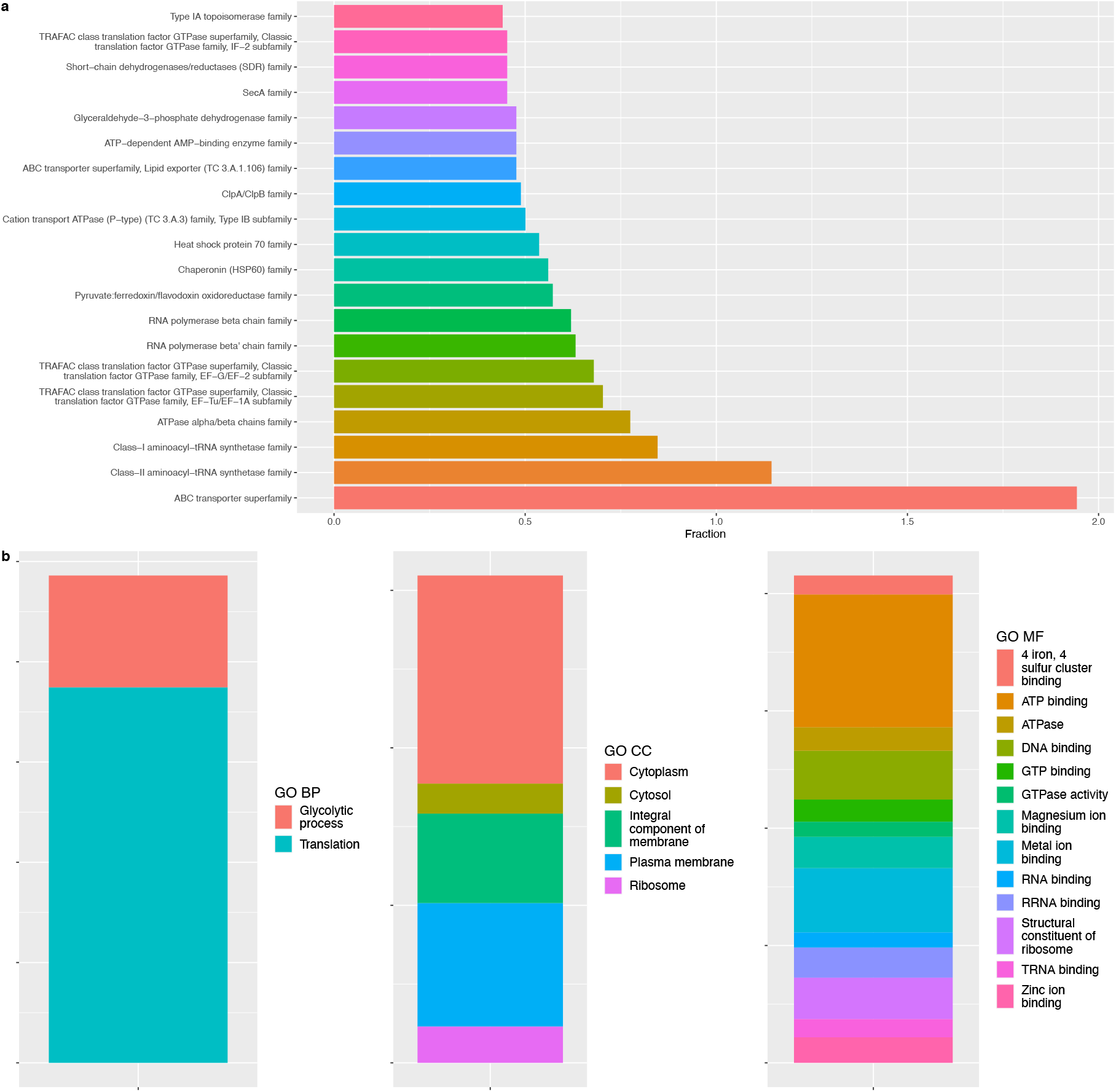
The most abundant (a) protein families and (b) Gene Ontology (GO) categories in all infected root canals.

As an additional functional annotation, we screened the three Gene Ontology GO databases: biological process (GO BP), molecular function (GO MF) and cellular component (GO CC). In total, 2,372 GO categories were associated to the 6,514 Uniprot ID (we remarked that a protein ID often belonged to multiple categories). Most of the identified GO terms belonged to GO MF (1,540), followed by GO BP (1,030) and GO CC (153). The 20 most abundant GO terms are shown in Figure 5b and included the major components of the cell (membrane, cytoplasm, ribosome), metal ion binding functions as well as ATP, GTP and RNA binding. Interestingly, among the 20 most abundant GO terms, GO CC were overrepresented by a factor of 3.5 and GO BP were underrepresented by a factor of 5 relative to the full set.

### 2.4. Differentially Expressed Genes

In order to identify genes whose expression was affected by the primary or secondary infection status, we tested transcripts for differential expression. Of the transcriptionally active 152 species, 76 were suitable to perform a differential expression analysis (see methods). We determined a total of 101 differentially expressed putative transcripts across 17 species. After removing the unannotated transcripts, we retained 11 species carrying 91 annotated Differentially Expressed Genes (DEGs), corresponding to 72 unique gene IDs [Supp. Table 6]. More than 85% of those DEGs were found in the three species harvesting the largest number of DEGs: *Cutibacterium acnes* (40), *Fusobacterium nucleatum* (21) and *Fusobacterium necrophorum* (17) [Figure 6a]. *Olsenella uli* is the only species among the 7 significantly more abundant in which we also found DEGs. A total of 17 genes were differentially expressed in more than one species, with ribosomal proteins *rpsL* and *rplR* common to three species, including the two aforementioned *Fusobacterium* spp. [Supp. Table 7]. The most represented protein families among DEGs were ribosomal protein families, which constituted about 30% of the DEG-associated families (27/91) and over 40% (7/17) of the DEG-associated families occurring in more than ones species [Figure 6b]. Interestingly, 23 of those 27 ribosome-associated DEGs were found either in *F. necrophorum* or *F. nucleatum*. We also report that only 3 DEGs, all in *C. acnes*, belonged to the ABC transporter superfamily, despite such protein family is the most abundant metatranscriptome-wide.

**Figure 6.**
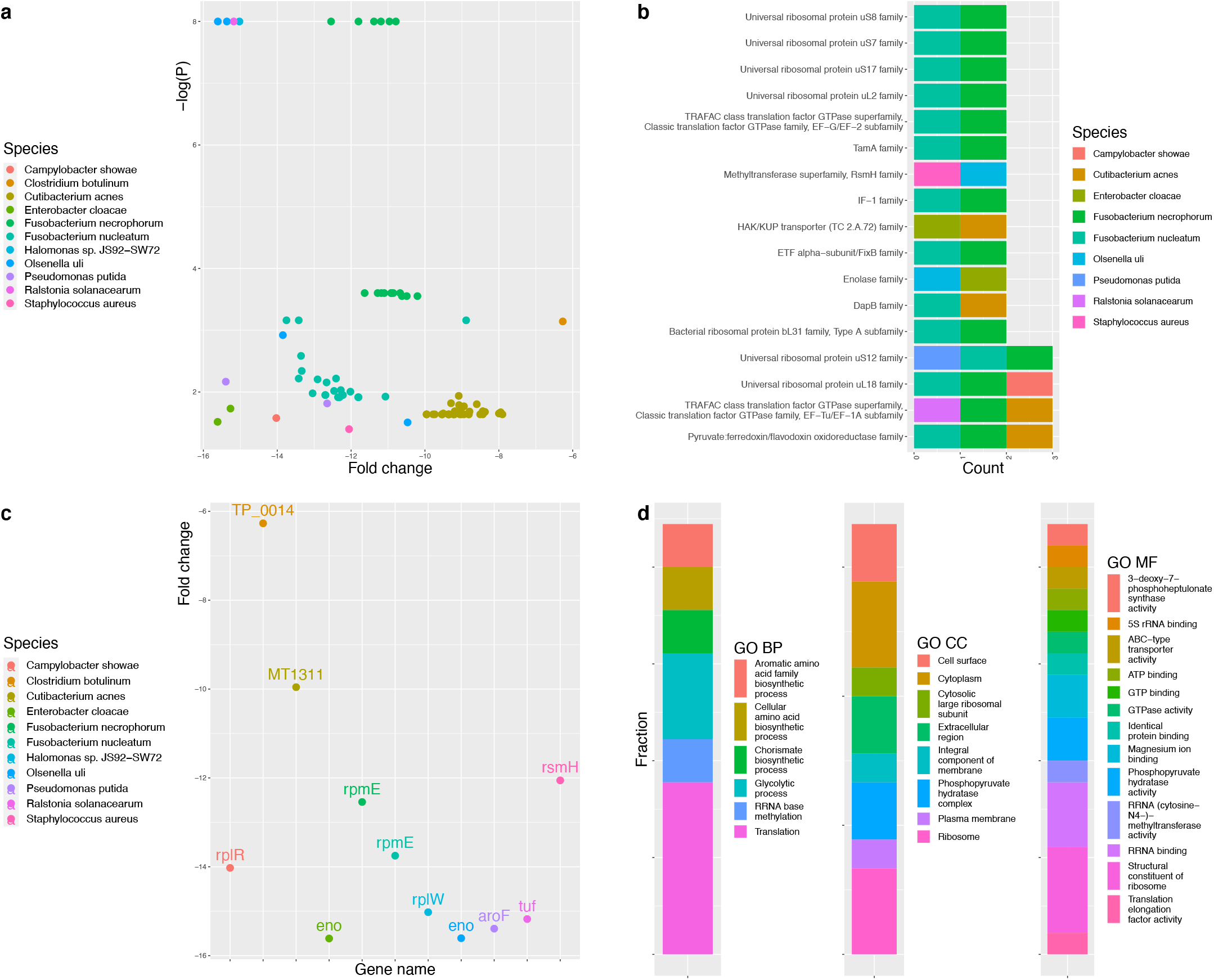
(a) Species with Differentially Expressed Genes (DEGs) in primary and secondary endodontic infections; (b) the most abundant protein families among DEGs; (c) the genes with the highest log-fold change to each species; and (d) GO categories associated with the 11 most downregulated genes.

The genes with the highest log-fold change to each species are shown in Figure 6c. Two genes, *eno* (Enolase) and *rpmE* (50S ribosomal protein L31) were the most downregulated between the groups in more than one species. Finally, the distribution of the GO terms associated with the 11 most downregulated genes are depicted in Figure 6d.

### 2.5. Microorganism-specific gene-set based functional analysis

The number of differentially expressed genes between the groups was rather low, therefore their potential functional and implications could be directly evaluated at the level of individual pathway and protein families. Nevertheless, for the three species with the largest numbers of DEGs, and for the species-wide pooled set of DEGs, we performed a gene-set based analysis. We interrogated the three Gene Ontology GO databases (GO BP, GO MF, GO CC), the KEGG pathway database and the INTERPRO protein domain databases with three lists of genes: those differentially expressed in *C. acnes* (40 unique Uniprot IDs), those differentially expressed in *F. nucleatum* and *F. necrophorum* (pooled, 26 unique Uniprot IDs) and the pool of all the annotated DEGs from 11 species (72 unique Uniprot IDs). In all cases, we used as background the list of 6514 unique Uniprot IDs that were identified metatranscriptome-wide.

No categories from any of the 5 databases resulted significantly overrepresented in the case of *C. acnes*. The DEGs from *Fusobacterium* spp. resulted in the enrichment of the GO BP category *translation* (p = 4e-09), GO CC categories *cytosolic large ribosomal subunits* (p = 9.3e-10) and *cytosolic small ribosomal subunits*, p= 3e-05), and GO MF categories *structural constituent of ribosome* (5.6e-10) and *rRNA binding* (4.4e-06). The significantly enriched categories obtained by pooling all the 72 DEGs were the same as those enriched when using the list from *Fusobacterium* spp., with slightly weaker p-values. These results are not surprising, as they confirm the observations made based on the individual gene annotations.

### 2.6. Antimicrobial resistance genes

Commensals in the oral microbiome regularly harbor a number of antimicrobial resistance (AMR) genes, so we found interesting to scrutinize the 8 root canal microbiomes for AMR genes. We mapped the reconstructed transcriptomes of the 152 species to the Comprehensive Antibiotic Resistance Database (CARD) of antibiotic resistance genes and we found a match in the case of 85 species, for a total of 423 genes homologs to 142 AMR gene IDs (as in the case of Uniprot, the match is at the level of the protein sequence, which in CARD can be annotated to a different species because the query species is not included in the database). More than half of the species (49/85) match only one or two genes, while for 8 species we report 10 or more homologs to AMR genes (Supp. Table 8). The 20 species with the largest number of AMR homologs are shown in Figure 7. The species with the largest number of AMR homologs was *Cutibacterium acnes*, followed by *Capnocytophaga* sp. oral taxon 323, *Fusobacterium nucleatum, Olsenella uli, Parvimonas micra, Prevotella intermedia, Staphylococcus epidermidis*, and *Tannerella forsythia*. Of interest, three differentially expressed genes from three species have homologs in CARD: *fusA*, a ribosomal elongation factor in both *F. nucleatum* and *F. necrophorum* matches to *tetB(P)*, a ribosomal protection protein conferring resistance to tetracycline; *MT1311* and *SDY_1145*, two ABC transporters in *C. acnes*, are homologs to the CARD entries *efrA* and *oleC*, respectively, both conferring antibiotic resistance via ABC transporters-mediated directed pumping of antibiotic out of a cell.

**Figure 7.**
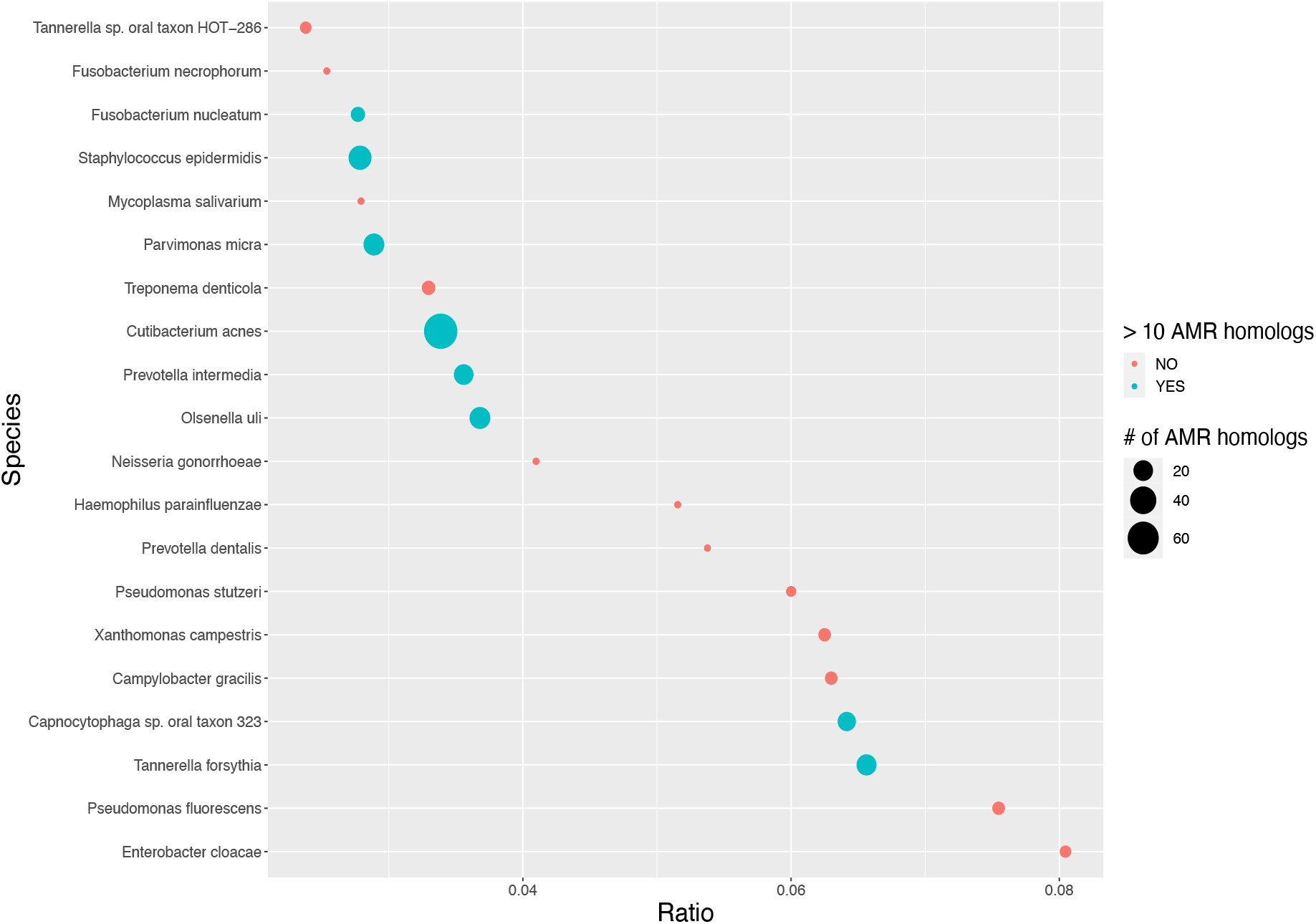
The 20 species with the largest number of AMR homologs. The species are sorted according to the increasing ratio between the number of AMR genes and total genes. The blue dots correspond to species with at least 10 AMR homologues. They are also listed in Table 2.

## 3. Discussion

The goal of this study was to explore the composition as well as the transcriptional and functional signatures of root canal microbiomes in teeth with asymptomatic apical periodontitis, including both primary and secondary endodontic infections (*i.e*., teeth with primary apical periodontitis or post-treatment apical periodontitis, respectively). We performed next-generation, microbiome-wide RNA-sequencing, generating, to the best of our knowledge, the first dataset capturing the metatranscriptome of infected root canals. The data indicated the presence of a core microbiome dominated by Proteobacteria, Bacteroidetes, Firmicutes and Actinobacteria, which is in agreement with previous reports on the endodontic microbiome [4-12]. The abundance of these phyla varied greatly between samples [6-12], reflecting a very dynamic and competitive ecosystem certainly shaped by individual conditions of the host environment. Among the non-dominant phyla, Fusobacteria, Spirochaetes and Synergistetes were observed in most samples. Despite being not-so-ubiquitous, these phyla have been identified in numerous other studies of the microbial populations in infected root canals [4-12], suggesting that they might represent a sub-core microbiome thriving in the peculiar conditions of an endodontic infection.

Metatranscriptomics data confirmed the activity of potential endodontic pathogens [2]. The top ten species were mainly represented by obligate (or quasi-obligate) anaerobes, including Gram-negatives (*e.g*., *Capnocytophaga* sp. oral taxon 323, *Fusobacterium nucleatum, Prevotella intermedia, Prevotella oris, Tannerella forsythia* and *Tannerella* sp. oral taxon HOT−286) and Gram-positive species (*e.g*., *Olsenella uli* and *Parvimonas micra*). In particular, *Prevotella* spp. and *O. uli* were more abundant in primary than in secondary infections. These findings are in agreement with previous reports showing that some species may be differentially abundant in primary and secondary endodontic infections [9,10]. This fact may be due to the environmental conditions in teeth with pulp necrosis that favor the growth of strict anaerobes capable of fermenting amino acids/peptides from necrotic pulp tissues and periapical fluids [22]. In turn, the microbial composition can be affected by ecological changes in the root canals after treatment, resulting in persistent infections (*i.e*., survival of bacteria from primary infections) or secondary infections (*i.e*., invasion of oral microorganisms via coronary microleakage). In the present study, *Bacillus cereus* was more abundant in secondary infections. Considering that *B. cereus* is not a normal inhabitant of primary endodontic infections, saliva may be the likely source of this opportunistic pathogen in cases of secondary infections [23]. In contrast to our findings, other authors found no significant differences between primary and secondary endodontic infections [6,8]. Discrepancies between studies may be due to differences in inclusion/exclusion criteria (*e.g*., the quality of previous endodontic treatment and the quality of coronal restorations), root canal sampling protocols, sequencing depth, and sample size.

Despite the limited number of root canal samples in the present study, the high sequencing depth allowed the expression analysis of a large set of genes (a total of 10,928 annotated transcripts in eight samples). *Streptococcus salivarius* showed the highest degree of transcriptional activity in the whole transcriptome. Gene related to histidine biosynthesis process (*hisB*) was one of the most abundant transcripts in this species. Similarly, histidine biosynthesis was overrepresented in cases of periodontal diseases [19]. Histidine catabolism can be an important source of carbon or nitrogen for other members of bacterial communities and may contribute to disease progression [16]. In addition, components of this pathway can affect bacterial properties such as virulence and biofilm formation. This may be particularly important for the pathogenesis of apical periodontitis, which, like other oral diseases, is a biofilm-induced disease [24].

Similar to a previous report on dental plaque biofilms, *gap* was the most expressed gene [17]. This gene encodes the enzyme GAPDH (glyceraldehyde-3-phosphate dehydrogenase) used in the process of glycolysis, a pathway commonly used in the metabolism of anaerobic bacteria. Although GAPDH is usually found in the bacterial cytosol, it can also appear on the cell surface or as a secreted protein, playing additional roles in bacterial interaction with the host [25]. The non-enzymatic functions of GAPDH have been shown to play important roles in the pathogenic processes of many bacteria, including streptococci and staphylococci [26]. For instance, GAPDH can bind to various human proteins, facilitate host colonization, and evade the host’s immune cells. Such proteins presenting two or more independent functions are referred to as moonlighting proteins [26]. Transcripts related to other moonlight proteins, such as pyruvate kinase (*pyk*), were also highly expressed in root canal samples. Pyruvate kinase participates in the last step of glycolysis leading to the production of ATP and pyruvic acid. In turn, this glycolytic enzyme can also act as an adhesin for yeast cells [26].

Most bacterial species expressed genes belonging to the same family, with ATP-binding cassette (ABC) transporters being the most predominant family. These findings are in agreement with previous reports of oral biofilms, indicating a high functional redundancy [14,15]. ABC transporters import essential nutrients for bacterial viability, including amino acids, peptides and metals [27,28]. This importer system facilitates bacterial adaptation to changes in the host environment. For example, the acquisition of peptides may favor the growth of pathogens commonly found in periodontal as well as endodontic infections (*e.g*., *P. micra, F. nucleatum* and *P. gingivalis*) [15]. They may also contribute to bacterial virulence through the export of virulent determinants such as toxins and proteins [27, 28]. Likewise, this system can export toxic xenobiotic substances, contributing to the development of antimicrobial resistance [27, 28]. In the present study, antibiotic resistance mediated by ABC transporters was detected in *Cutibacterium acnes*.

Other functional signatures include those involved in protein translation and energy metabolism, which is similar to a previous report on dental plaque transcriptome [17]. The aminoacyl-tRNA synthetase families, one of the most abundant in the studied samples, play a key role in protein synthesis by pairing tRNAs with amino acids [29]. An interesting fact is that they have been considered as potential targets for the development of new antimicrobial agents, which may be of clinical interest in dentistry [30]. Likewise, the GTPase families, which encode the translational elongation factors EF-Tu and EF-G, appear among the most abundant transcripts. Translational elongation factors are also moonlight proteins, which, in addition to participating in protein synthesis, may favor bacterial adhesion [26, 31].

Transcripts encoding stress response proteins were also abundant in root canal samples, which probably contributes to bacterial adaptation in this environment [32]. For instance, the pyruvate:ferredoxin/flavodoxin oxidoreductase family may contribute to bacterial adaptation to oxidative stress, while the chaperonin (HSP60) and heat shock protein 70 families may promote protein recovery from stress-induced unfolding [33]. Interestingly, chaperonin and heat shock proteins are also moonlighting proteins involved in bacterial adhesion and biofilm formation [26]. In addition, chaperonin can synergize with LPS from Gram-negative bacteria, activate cytokine synthesis and stimulate bone resorption [26], which are key factors in the development of apical periodontitis.

*Fusobacterium* species were the main players in primary endodontic infections. Despite not being abundant, *F. nucleatum* and *F. necrophorum* showed a large number of DEGs (Differentially Expressed Genes). These findings are in agreement with a previous transcriptomic analysis of periodontal samples, showing that although the proportions of *F. nucleatum* did not change between health and disease, bacterial metabolism was different in periodontitis samples [16]. Therefore, *F. nucleatum* has been proposed as a keystone species for the development of periodontal disease [16]. In endodontics, *F. nucleatum* has been associated with primary infections [10], especially in cases with clinical symptoms [34]. As the present study included only asymptomatic cases, future transcriptomic analysis should investigate the metabolism of *Fusobacterium* spp. in cases of acute endodontic infections. In the present study, most transcripts from *Fusobacterium* spp. were involved in protein synthesis, represented mainly by the Ribosomal protein-, GTPase-, and IF-1 families. The gene-set based functional analysis highlighted the enrichment of the GO BP category *translation*; GO CC categories *cytosolic large and small ribosomal subunits*; and GO MF categories *structural constituent of ribosome* and *rRNA binding*. In addition to transcripts involved in protein synthesis, *Fusobacterium* spp. also produced transcripts encoding oxidative stress responses (*e.g*., pyruvate:ferredoxin/ flavodoxin oxidoreductase family). Similar to findings from other dental biofilms [17], these data suggest that oxidative stress may be one of the main stressors in endodontic biofilms.

*Cutibacterium acnes* (formerly known as *Propionibacterium acnes*) was the species with the highest number of AMR homologs (71). This fact may explain the resistance of this species to antimicrobial procedures commonly used during endodontic treatment [35]. *C. acnes* can persist active in root canals after root canal preparation, being a possible source for the establishment of persistent infections [36]. Other bacterial species commonly found in endodontic infections (e.g., *Capnocytophaga* sp., *F. nucleatum, O. uli, P. micra, P. intermedia* and *T. forsythia*) expressed high numbers of AMR homologs. In particular, *tet*B-like genes were overexpressed by *Fusubacterium* spp. These findings are in agreement with a previous systematic review showing that endodontic isolates have a high rate of tetracycline resistance [37]. More importantly, these findings highlight that oral species may represent a reservoir of resistance to several antibiotics [38].

## 4. Conclusions

Metatranscriptomics data indicated the presence of a core microbiome in root canals dominated by Proteobacteria, Bacteroidetes, Firmicutes and Actinobacteria. Among the non-dominant phyla, Fusobacteria, Spirochaetes and Synergistetes were observed in most samples. At species level, the microbiome was dominated by Gram-negative and Gram-positive anaerobes. In particular, *Prevotella* spp. and *O. uli* were more abundant in primary than in secondary infections. An interesting finding of the present study was the abundance of transcripts encoding moonlighting proteins in infected root canals, including glycolytic proteins (*e.g*., glyceraldehyde-3-phosphate dehydrogenase and pyruvate kinase), translational elongation factors (*e.g*., EF-Tu and EF-G), chaperonin and heat shock proteins. Considering their potential effect on bacterial virulence (*e.g*., biofilm formation, evasion of host defense, and induction of inflammation), future studies should investigate the possible role of moonlighting proteins in the pathogenesis of apical periodontitis. Functional analysis revealed that *Fusobacterium* species were the main players in primary endodontic infections, although they were not dominant members. *Fusobacterium* transcripts were involved in protein synthesis, oxidative stress responses and antibiotic resistance, especially to tetracycline. Moreover, other endodontic pathogens expressed high numbers of AMR homologs.

## 5. Materials and Methods

### 5.1. Ethics statement

This study was conducted in accordance with the Declaration of Helsinki and approved by the Ethics Committee of the Faculty of Dentistry of the University of São Paulo (nº 2,201,768; 08/04/2017). All participants signed an informed consent form before treatment.

### 5.2. Patient enrollment and clinical diagnosis

Adult patients (18-65 years) who required endodontic treatment/retreatment of single-rooted teeth with apical periodontitis were included in this study. Evidence of apical periodontitis was determined by radiographic examination showing bone destruction around the root apex of the tooth, which is a consequence of root canal infections in teeth with necrotic pulps (primary infections) or teeth that have been previously root-filled (persistent/ secondary infections). Endodontic treatment failure was determined by the presence of apical periodontitis in teeth treated for more than 4 years, indicating the absence of apical repair and the need for endodontic retreatment. Although most teeth included in this study were not intact (*i.e*., there was caries or restorations that clinically/ radiographically appeared to be unsealed), teeth with great loss of coronary structure that could not be adequately isolated with a rubber were excluded. In addition, patients who received antibiotic therapy during the previous 3 months or with periodontal pockets depths greater than 4 mm were excluded from the study.

### 5.3. Root canal samples

Root canal samples were collected from ten patients, five with primary infections and five with persistent/secondary endodontic infections (called secondary infections in this study). Sampling procedures were performed as previously described [12,36,39]. After removal of dental plaque and caries, the tooth and rubber dam were disinfected with 30% H_2_O_2_, 2.5% NaOCl and 5% sodium thiosulfate. The access cavity was performed under irrigation with sterile saline solution. Before entering the pulp chamber, the disinfection protocol was repeated and the access cavity completed. Then, a control sample from the disinfected cavity was performed with paper cones and stored at −80°C. The absence of bacteria in the control samples was verified by PCR and RT-PCR using universal primers for Bacteria.

In primary endodontic infections, the root canal was filled with sterile saline and the working length was established using an electronic apex locator (J. Morita Brasil, São Paulo, SP, Brazil). After applying a filing motion with a Hedström file, five sterile paper points were placed consecutively at working length (1 minute each) and placed in cryotubes containing 300 μL of RNAlater solution, which were stored at −80°C. In cases of secondary endodontic infections, the root filling was removed with reciprocating instruments (Wave One file, Dentsply Sirona, Ballaigues, Switzerland) and irrigation with sterile saline solution. Then, root canal sampling was performed as described above.

### 5.4. RNA Extraction

RNA from endodontic samples was extracted using the MasterPure Complete DNA and RNA Purification Kit (Epicentre Technologies, Madison, WI, USA) following the manufacturer’s protocol. Prior to reverse transcription, an additional treatment with DNase I (ThermoFisher Scientific, MA, USA) was performed and the absence of contaminating DNA was confirmed by PCR using universal primers. Then, complementary DNA (cDNA) synthesis was performed using the SuperScript III First-Strand Synthesis System (ThermoFisher Scientific, MA, USA) and quantified using a Qubit fluorometer (ThermoFisher Scientific, MA, USA). The amount of material in two samples from secondary infections was insufficient to measure microbial gene expression, reducing the number of samples to eight (five from primary infections and three from secondary infections). RNA sequencing was performed on a NovaSeq6000 system (Illumina, San Diego, CA, USA).

### 5.5. Metatranscriptomics analysis

The raw reads were preprocessed using Trimmomatic v0.39 [40] with the parameters “*ILLUMINACLIP:adapters.fa:1:30:10 LEADING:0 TRAILING:0 SLIDINGWINDOW:5:15 AVGQUAL:20 MINLEN:80*”. We then assembled the preprocessed reads using Trinity v2.11 [41] with default settings. We used Transdecoder v5.5.0 [42] to reduce the redundancy of the resulting set of putative transcripts and mapped the reduced set of sequences against the kraken2 bacterial database [43]. The sequences with a taxonomic classification were then annotated against Uniprot using diamond v2.0.5 [44] with parameters “*-k 10 -e 0.001”*. Set-based gene analysis was performed using DAVID [45]. Finally, we identified differentially expressed genes using a genewise binomial linear model and quasi-likelihood test as implemented in the functions *glmQLFit* and *glmQLFitTest* in the R Bioconductor package EdgeR [46]. We adjusted the resulting p-values for multiples testing using the Benjamini-Hochberg procedure.

## Supplementary Materials

The following supporting information can be downloaded at: www.mdpi.com/xxx/s1, Figure S1: title; Table S1: title; Video S1: title.

## Author Contributions

Conceptualization, E.T.P., L.K., and T.T.; formal analysis, G.R.; investigation, E.T.P., G.T.M.C., and B.G.V.; writing— original draft preparation, E.T.P. and G.R.; writing—review and editing, B.P.F.A.G., T.A., L.K., and T.T.; funding acquisition, E.T.P., L.K., and T.T. All authors have read and agreed to the published version of the manuscript.

## Funding

This research was funded by São Paulo Research Foundation (FAPESP 2019/12908-3; 2021/14434-9).

## Institutional Review Board Statement

This study was conducted in accordance with the Declaration of Helsinki and approved by the Ethics Committee of the Faculty of Dentistry of the University of São Paulo (nº 2,201,768; 08/04/2017).

## Informed Consent Statement

Informed consent was obtained from all subjects involved in the study.

## Data Availability Statement

the preprocessed are being submitted to the NCBI sequence read archive and the final submission number will be available prior to the publication.

## Acknowledgments

We would like to thank the Functional Genomics Center Zurich (FGCZ) at the University of Zurich for providing the sequencing services.

## Conflicts of Interest

The authors declare no conflict of interest.

## Notes

### Competing Interest Statement

The authors have declared no competing interest.

